# Interactions between the Bone Morphogenetic Protein and the Planar Cell Polarity Pathways lead to distinctive ethanol-induced facial defects

**DOI:** 10.1101/2025.04.23.650288

**Authors:** Raèden Gray, Anna Llyod, C. Ben Lovely

**Affiliations:** University of Louisville, School of Medicine, Department of Biochemistry and Molecular Genetics and Alcohol Research Center, Louisville, KY

**Keywords:** FASD, zebrafish, genetics, craniofacial development

## Abstract

**Background:** Fetal Alcohol Spectrum Disorders (FASD) describes a spectrum of ethanol-induced neural and facial developmental defects. Ethanol susceptibility is modulated by genetics, but their underlying mechanisms remain poorly understood. In all vertebrates, a series of complex cellular events give rise to the body plan, including convergence & extension (C&E) and endoderm/ cranial neural crest (CNC-which gives rise to the facial skeleton) morphogenesis. These events are critical to establish complex signaling interactions, driving embryo development, including the facial skeleton. In zebrafish, C&E occurs between 6-10 hpf while endoderm/CNC morphogenesis occurs 10-24 hpf. Previous work shows that the PCP mutants are sensitive to ethanol from 6-24 hpf, covering both C&E and endoderm/CNC morphogenesis and exhibiting multiple defects to the forming head raising the question whether ethanol during both time windows drives PCP-ethanol defects. We hypothesize that PCP single and double mutants are ethanol sensitive 10-24 hpf, after C&E. We also hypothesize BMP signaling (sensitive 10-18 hpf) interacts with and sensitizes the PCP pathway to ethanol.

**Methods:** Here, we treated PCP/BMP mutants with ethanol from 6-10, 10-18, 10-24 or 24-30 hpf and combined morphometric and linear measurements to examine facial development.

**Results:** We show that PCP mutant larvae are ethanol-sensitive from 10-24 hpf, but not 6-10 or 24-30 hpf. We also show that BMP mutants sensitize PCP mutants to ethanol and lead to novel ethanol-independent midline craniofacial defects. Our results suggest that the ethanol-sensitive role of PCP pathway occurs after C&E, during endoderm/CNC morphogenesis and that the PCP and BMP pathways genetically interact during the morphogenesis events.

**Conclusions:** Ultimately, our work builds on a mechanistic paradigm of ethanol-induced birth defects we have been developing, connecting conceptual framework with concrete cellular events that could be ethanol-sensitive beyond facial development.

## Introduction

Fetal Alcohol Spectrum Disorders (FASD) describes a wide range of birth defects due to prenatal alcohol exposure (PAE) (Sokol et al. 2003; Hoyme et al. 2016). Although PAE is the most preventable cause of birth defects, FASD still impacts up to 5% of children in the US, with estimates even higher in other regions across the globe (May et al., 2018; Popova et al., 2020). However, these are likely underestimates as up to 10% of women worldwide consume alcohol during pregnancy, though some estimates are higher such as 23.3% of women in the Caribbean, for example (Popova et al., 2018, 2017; Landgraf et al., 2013; Lange et al., 2017). In addition, up to 50% of pregnancies in the US are unplanned and many pediatricians fail to recognize and diagnose FASD (Rojmahamongkol et al., 2015; Finer and Zolna, 2016). Overall, PAE is one, if not the leading cause of birth defects.

Fetal Alcohol Syndrome (FAS) is the most severe form of FASD and is characterized by neural, growth and craniofacial defects (Landgraf et al., 2013; May et al., 2009). Not only can craniofacial defects negatively impact the ability for children to eat and drink (Perkins et al., 1997), sociologists have shown that children who have craniofacial defects, such as cleft lip/palate, face a significant amount of stigma and social rejection (Chung et al., 2019), increasing disadvantages the child will face in social environments. These factors together explain why the cost of correcting these and other congenital birth defects are over a billion dollars annually (Swanson, 2023). While the developmental window of exposure and dosage of ethanol contribute to the etiology of FASD, human twin studies demonstrate that genetics also modulate the effects of prenatal ethanol exposure, where there is 100% concordance in FASD phenotypic outcomes in monozygotic twins over dizygotic twins (66%), full siblings (41%) and half siblings (22%) (Hemingway et al., 2019). However, the ethanol-sensitive genetic loci that contribute to FAS susceptibility are still not well understood. The social disadvantages as well as the physical impact of the craniofacial defects caused by PAE highlights how imperative it is to understand how genetic and environmental factors intersect, driving craniofacial malformations observed in FASD.

A key phenotype of FASD is jaw hypoplasia or reduction in jaw size. The jaw forms from the mandibular domain of the cranial neural crest (CNC) in the first pharyngeal arch (Chai et al., 2000). CNC development into the jaw is dependent on the presence of the pharyngeal endoderm. Previous studies show that the pharyngeal endoderm acts as a signaling center through tissue-tissue interactions that provide extracellular cues to the CNC, oral ectoderm, and mesoderm (Chai et al., 2006). In embryos that lack endoderm the CNC condense into the arches but undergo apoptosis and fail to form the facial skeleton (Trumpp et al., 1999; David et al., 2002). Many genes involved in CNC morphogenesis of the jaw are not active until after the mandibular domain has already condensed, showing jaw development’s reliance on these cues (Trumpp et al. 1999). Additional studies have shown that when the anterior pharyngeal endoderm is malformed, as seen in zebrafish mutants in Sphingosine-1-phosphate (S1P) signaling, it leads to jaw malformation (Balczerski et al., 2012). Together, this emphasizes the importance of proper pharyngeal endoderm morphogenesis for development of the jaw.

Multiple signaling pathways are critical for endoderm morphogenesis and jaw development, including S1P signaling (described above), Fibroblast Growth Factor (Fgf) signaling and Bone Morphogenetic Protein (BMP) Signaling (Lovely et al., 2016). We and others have shown that the BMP pathway is critical for morphogenesis of the endoderm by regulating a series of downstream signaling targets regulating endodermal cell behaviors (Lovely et al., 2016; Li et al., 2019). We have shown that when BMP signaling is blocked with the small chemical inhibitor Dorsomorphin (DM) from 10-18 hours post fertilization (hpf) in zebrafish embryos, overall endoderm morphogenesis is disrupted leading to a wide range of craniofacial malformations (Lovely et al., 2016). We have recently shown that mutations in multiple genes of the BMP pathway, such as *bmp4*, sensitize embryos to ethanol-induced defects to the anterior endoderm leading to defects in both the jaw and palate (Lovely, C.B., 2024; Klem et al., 2025). However, we show that ethanol does not disrupt BMP signaling directly, nor does it reduce expression of the downstream BMP target, *nxk2.3* (Klem et al., 2025, Vo and Lovely, 2025). In addition, knockdown of *nxk2.3* does not sensitive larvae to ethanol-induced facial defects (Vo and Lovely, 2025). This suggests that ethanol is acting on additional loci, contributing to the facial defects in ethanol-treated BMP mutants.

Previous work has shown that components of the Planar Cell Polarity (PCP) pathway are ethanol sensitive resulting in a range of phenotypes including craniofacial defects (Sidik et al., 2021; Swartz et al., 2014). Mutants in *vangl2* and *gpc4* are ethanol sensitive leading to disrupted axonal projections, synophthalmia and a range of defects to the facial skeleton, which were exacerbated in the double mutant embryos (Swartz et al., 2014; Sidik et al., 2021). Work from Sidik et al (2021) showed that *vangl2* mutants are sensitive to ethanol leading from 6-24 hpf, covering both early convergence & extension and subsequent endoderm morphogenesis. Convergence & extension (C&E) describes a crucial part of embryogenesis in which germ layers elongate and narrow to develop the animal body plan (Keller, 2002) and typically occurs in zebrafish between 6 and 10 hpf (Warga & Kimmel, 1990; Topczewski et al., 2001; Jessen et al., 2002; Sepich et al., 2005;). The PCP pathway drives C&E by modulating the cell movement by regulating cell polarity and cell adhesion (Muñoz-Soriano et al., 2012). Vangl2 is a transmembrane protein in the pathway whose phosphorylation is induced by Wnt ligands and is needed to promote PCP signaling and, in part, regulate E-cadherin levels and distribution (Dush & Nascone-Yoder,2019; Yang et al., 2017). Gpc4 is a cell surface proteoglycan required for the transport of Wnt ligand and has been shown to be important in Wnt ligand transport from endoderm to other germ layers (Hu et al., 2021). In addition, previous work shows that the PCP components *vangl2* and *gpc4* interact and play a key role during early C&E of the endoderm (Miles et al., 2017). Noticeably, there were ethanol-induced jaw defects in both *vangl2* and *gpc4* mutants, suggesting that these ethanol-sensitive PCP mutants may play a role in craniofacial development (Sidik et al., 2021; Swartz et al., 2014). However, the timing of ethanol sensitivity of these jaw defects was not further examined raising the question of the timing of the ethanol sensitive role *vangl2* and *gpc4* play in facial development.

Here, we examine the role of *vangl2* and *gpc4,* and their interaction with BMP signaling, in jaw development, post C&E, and what ethanol may be doing to disrupt that role. We show that *vangl2* and *gpc4* are sensitive to ethanol-induced jaw defects starting at 10 hpf, after C&E, with *vangl2*; *gpc4* double mutants being hypersensitized. We go on to show that loss of *bmp4* further sensitizes *vangl2* or *gpc4* mutants to ethanol-induced craniofacial defects, resulting in novel ethanol-induced midline facial defects not previously observed in PCP or BMP single mutants alone. Collectively, our data links perturbations in PCP and BMP signaling to ethanol susceptibility during jaw development, establishing a genetic pathway in gene-ethanol interactions for future studies in FASD.

## Materials and Methods

### Zebrafish (Danio rerio) care and use

All zebrafish were raised and cared for using established IACUC protocols approved by the University of Louisville (Westerfield, 2007). Adult fish were maintained at 28.5°C with a 14 / 10-hour light / dark cycle. The *vangl2^m209^*(Solnica-Krezel et al., 1996; Stemple et al., 1996; Driever et al., 1996), *gpc4^fr6^* (Topczewski et al., 2001) and *bmp4^st72^* (Stickney et al., 2007) zebrafish lines used were previously described.

### Zebrafish staging and ethanol treatment

Embryos were collected, morphologically staged as previously described (Westerfield, 2007), sorted into sample groups and reared at 28.5°C in embryo media (EM) to desired developmental time points. At 6 hpf, 8 hpf, 10 hpf, and 24 hpf EM was changed to either fresh EM or EM containing 1% ethanol (v/v) depending on the time window being examined. At 10 hpf, 12 hpf, 18 hpf, 24 hpf and 30 hpf, EM containing ethanol was washed out with 3 fresh changes of EM.

### Alcian Blue cartilage staining

Zebrafish larvae were fixed at 5 dpf (single mutants) and 4 dpf (double mutants due to increased die off at 5 dpf) and the facial cartilages were stained with alcian blue (Walker & Kimmel, 2007). Whole mount, ventral view, brightfield images of the viscerocranium were taken on an Olympus BX53 compound microscope.

### Morphometric analyses

Morphometric analysis of Alcian-stained larva was performed in TpsDig2 (https://sbmorphomectrics.org) and MorphoJ (Klingenberg, 2011). Landmarks were placed on the following joints, Meckel’s cartilage midline joint, the joints between Meckel’s’ and the palatoquadrate, the palatoquadrate and ceratohyal and at the end of the hyomandibular cartilages. Linear measures were analyzed using TpsDig2. Principle component analysis (PCA), Procrustes ANOVA and wireframe graphs of facial variation were generated using MorphoJ.

### Statistical Analysis

Cartilage angles and linear measures of Alcian-stained viscerocranium were analyzed with a two-way ANOVA (type III) and Sidak’s multiple comparison test in Graphpad Prism 10.4.1 (Graphpad Software Inc., La Jolla, CA).

## Results

### Ethanol sensitizes *vangl2* and *gpc4* to ethanol induced jaw defects from 10 – 18 hpf

Previous work from Sidik et al., (2021) showed that when exposed to ethanol between 6-24 hpf, *vangl2* and *gpc4* zebrafish mutants exhibited a range of defects including those to the facial skeleton. This time window encompasses a host of developmental processes, including convergence & extension (C&E), somitogenesis, anterior pharyngeal endoderm (APE) morphogenesis, and cranial neural crest (CNC) migration and condensation. Because the role of CNC and APE in jaw development occur after C&E, we hypothesize that *vangl2* & *gpc4* are required for jaw development after 10 hpf. To test this, we exposed our wildtype and *vangl2* or *gpc4* mutant embryos to 1% ethanol from 10 hpf to 24 hpf. We selected 1% ethanol (v/v) as it is the highest dose that does not cause craniofacial defects in wild-type larvae, while higher doses have been shown to impact facial development (Bilotta et al., 2004; Everson et al., 2022; McCarthy et al., 2013; Swartz et al., 2014; Zhang et al., 2014). 1% ethanol equilibrates to approximately 30% of the media within 5 minutes of exposure (Flentke et al., 2014; Lovely et al., 2014; Reimers et al., 2004; Zhang et al., 2013), resulting in an embryonic ethanol concentration of 50 mM, roughly equivalent to a human Blood Alcohol Concentration of 0.23. While this is a binge dose, it is physiologically relevant to FASD with humans readily surpassing this amount (Canfield et al., 2019; Ethen et al., 2009; Jones, 2008; Maier, 2001; Whaley et al., 2019).

We stained the facial cartilage with Alcian blue and used morphometric analysis to observe facial shape changes between our untreated and ethanol-treated embryos. The benefit of utilizing morphometric analysis is that we can see the change in shape, independent of general size effects, which would not be identified using traditional linear measurements. We were able to generate a principal component analysis plot to visualize the variation between our groups. Our data show that untreated *vangl2* mutants have mild facial shape changes compared to untreated and ethanol-treated wild-type siblings (Figure 1A-C). This results in a shift in PC1, which denotes a shortening and widening of the viscerocranium and a flattening of the ceratohyal (Ch) and Meckel’s (Mc) cartilages and represents 80% of all variation in the data (Figure 1E). When exposed to ethanol from 10-24 hpf, *vangl2* mutants have increased variation in jaw shape relative to untreated mutants, shifting further along PC1 (Figure 1D compared to C, E). PC2, which denotes subtle shape change in the position and angle of both the Ch and (Figure 1A-C, E). PC2. represents only 4% of variation in the data (Figure 1E). This suggests that while untreated *vangl2* mutants have subtle facial shape changes not observed before ethanol exposure from 10-24 hpf, after C&E, leads to a significant change in facial shape.

**Figure 1.**
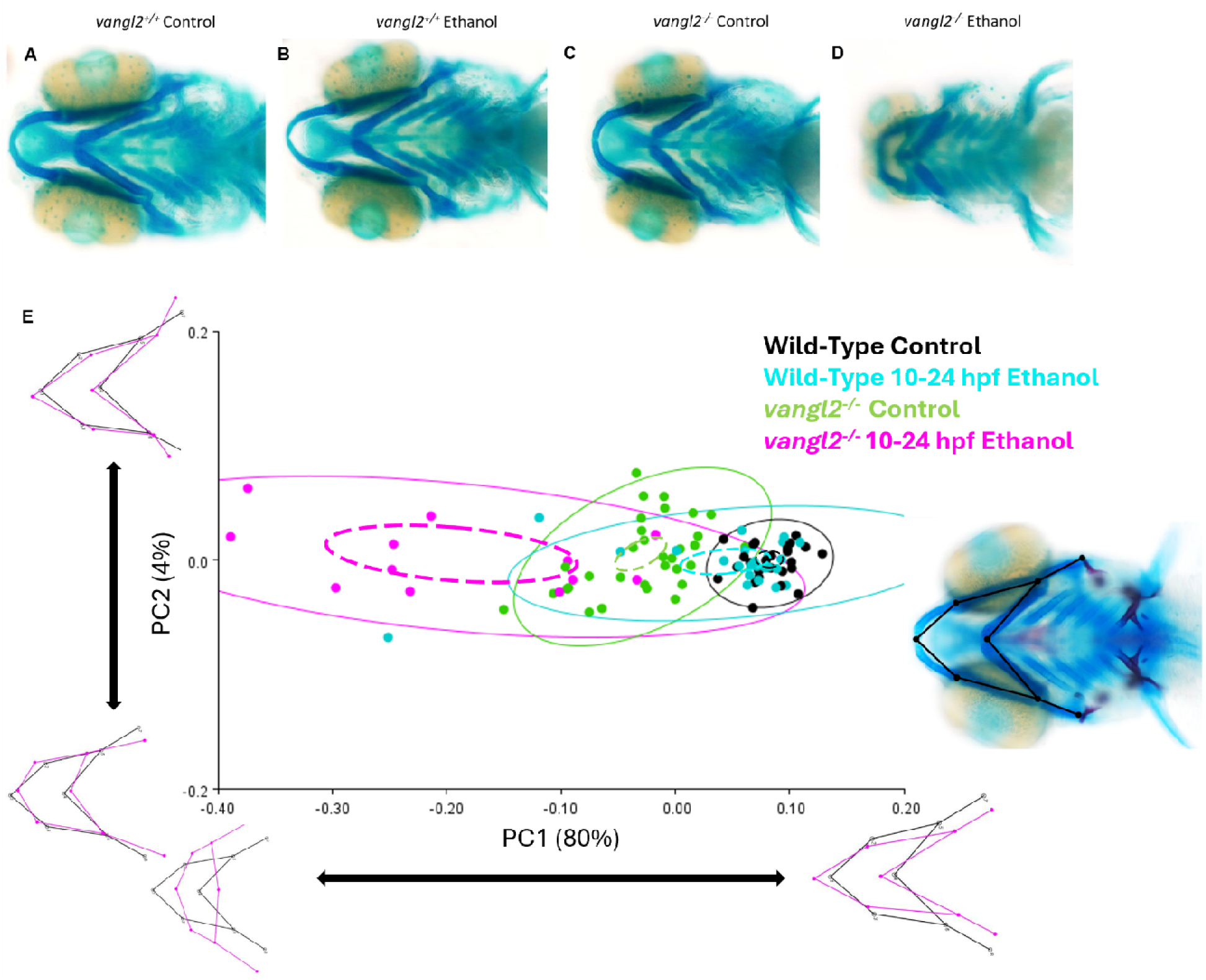
*vangl2* mutants are sensitized to ethanol induced jaw hypoplasia. Whole-mount images of the viscerocranium of **(A)** wild type control embryos (B) wild type ethanol treated embryos, **(C)** *vangl2^−/−^* control fish and **(D)** *vangl2^−/−^* ethanol treated fish. **(E)** Principal component analysis plot and wireframes from morphometric analysis data. Landmarks were placed on the joints between the cartilage structures of the viscerocranium. Each genotype is color-coded: black = wild type control embryos (n=25), cyan = wild type ethanol treated embryos (n=25), green n= *vangl2^−/−^* control embryos (n=32), magenta = *vangl2^−/−^* ethanol treated embryos (n=12), solid circles represent confidence ellipses in which 95% of all individual data points for each group lie, dashed circles represent 95% confidence ellipses for means. Wireframe graphs represent variation termed by each axis with black representing no variation and magenta representing variation relative to the black wireframe. For example, PC1 captures a shortening and widening in viscerocranial shape, while PC2 represents variation in midfacial width. Procrustes ANOVA showed significant change in the viscerocranial shape (F = 32.63, DF = 36, p = <.0001).

Previous studies have shown that *gpc4* mutants have craniofacial defects, including jaw hypoplasia, in control conditions (Sisson et al., 2015; LeClair et al., 2009). We observed similar facial defects in our morphometric approach, with untreated *gpc4* mutants having shorter and wider viscerocranium and flattened Ch and Mc cartilages (Figure 2A-C, E). Here, the shortening and widening of the viscerocranium in PC1 represents 86% of the variation in the data (Figure 2E). However, in contrast to ethanol-treated *vangl2* mutants, ethanol-treated *gpc4* mutants show little difference in facial shape compared to their untreated mutant siblings (Figure 2D compared to C, E). We observed an interesting shift in facial shape between untreated and ethanol-treated *gpc4* mutants. While untreated mutants were relatively consistent along PC1, they had greater spread over PC2 (representing 7% of the variation) which denotes a flattening of the Ch leading to greater distance between the midline joints of the Mc and the Ch (Figure 2E). On the other hand, ethanol-treated gpc4 mutants showed greater spread along PC1, which includes an inversion of the Ch with the shortening and widening of the viscerocranium (Figure 2E). Therefore, it appears that while more subtle in their ethanol-induced facial defects compared to *vangl2* mutants, *gpc4* mutants still display ethanol-induced defects in viscerocranial shape when treated from 10-24 hpf, after C&E.

**Figure 2.**
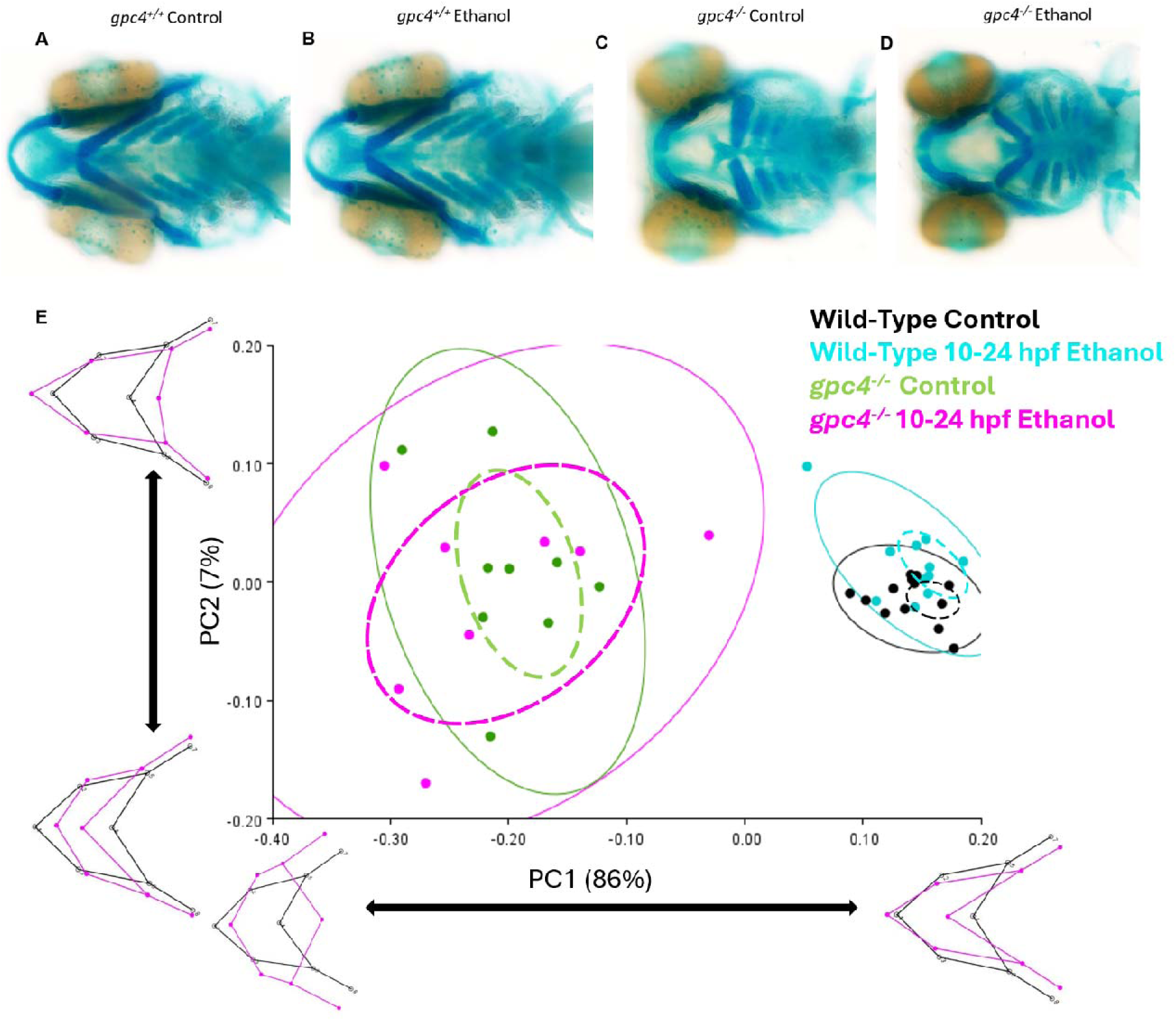
Ethanol drives greater variation in viscerocranial shape in *gpc4* mutant embryos. Whole-mount i ages of the viscerocranium of **(A)** wild type control embryos **(B)** wild type ethanol treated embryos, **(C)** *gpc4^−/−^* control fish and **(D)** *gpc4^−/−^* ethanol treated fish. **(E)** Principal component analysis plot and wireframes from morphometric analysis data. Landmarks were placed on the joints between the cartilage structures of the viscerocranium. Each genotype is color-coded: black = wild type control embryos (n=13), cyan = wild type ethanol treated embryos (n=12), green = *gpc4^−/−^* control embryos (n=9), magenta = *gpc4^−/−^*ethanol treated embryos (n=8), solid circles represent confidence ellipses in which 95% of all individual data points for ach group lie, dashed circles represent 95% confidence ellipses for means. Wireframe graphs represent variation termed by each axis with black representing no variation and magenta representing variation relative to the black wireframe. For example, PC1 captures a shortening and widening in viscerocranial shape, while C2 represents variation in midfacial width. Procrustes ANOVA showed significant change in the viscerocranial shape (F = 51.25, DF = 36, p = <.0001).

In addition to facial defects ethanol-treated *vangl2* mutants also showed eye field defects, where the majority of ethanol-treated *vangl2* mutants displayed synophthalmia (eye field fusions) with a smaller minority showing full cyclopia (complete lens fusion; Swartz et al., 2014; Sidik et al., 2021). However, these studies both exposed *vangl2* mutants to ethanol from 6-24 hpf, overlapping with C&E and Sidik et al, (2021) showed that failure of eye field separation was driven in part by disrupted C&E. In our exposure paradigm, we did observe synophthalmia in 50% of ethanol-treated *vangl2* mutants but not in our *gpc4* mutants (Figures 1D, 2D, Table 1). While phenotypically consistent with previous work (Swartz et al., 2014; Sidik et al., 2021), our exposure paradigm begins at 10 hpf, after C&E, suggesting that ethanol disrupts eye field separation independent of C&E, arguing other signaling mechanism may be disrupted.

**Table 1.** Mutations in *vangl2* and *gpc4* sensitize embryos to eye-field defects. A composite summary of the percentage of embryos that have specific eye-field phenotypes. The x-axis represents the genotypes (Wild-Type, *vangl2* mutant, *gpc4* mutant, *vangl2* mut *gpc4* het, *gpc4* mut *vangl2* het and double mutant) and treatment conditions (Control or Ethanol) evaluated. The y-axis represents the eye phenotypes: normal eye-field separation synophthalmia (eye fusion), synophthalmia (eye & lens fusion) and cyclopia.

|  | Wild-Type |  | *vangl2* mut |  | *gpc4* mut |  | *vangl2* mut *gpc4* het |  | *gpc4* mut *vangl2* het |  | double mutant |  |
| --- | --- | --- | --- | --- | --- | --- | --- | --- | --- | --- | --- | --- |
|  | Control | Ethanol | Control | Ethanol | Control | Ethanol | Control | Ethanol | Control | Ethanol | Control | Ethanol |
| Normal Eye-field Separation | 100% | 100% | 100% | 50% | 100% | 100% | 88% | 59% | 88% | 13% | 0% | 0% |
| Synophthalmia (Eye Fusion) | 0% | 0% | 0% | 50% | 0% | 0% | 12% | 41% | 12% | 83% | 80% | 27% |
| Synophthalmia (Eye & Lens Fusion) | 0% | 0% | 0% | 0% | 0% | 0% | 0% | 0% | 0% | 0% | 0% | 64% |
| Cyclopia | 0% | 0% | 0% | 0% | 0% | 0% | 0% | 0% | 0% | 4% | 20% | 9% |

To further confirm that both *vangl2* and *gpc4* are ethanol sensitive after C&E and during APE and CNC morphogenesis, we explored the following multiple ethanol exposure time windows: 1. during C&E (6-10 hpf); 2. when we have previously shown that BMP signaling is required for APE morphogenesis (10-18 hpf); 3. after CNC condensation into the pharyngeal arches (24-30 hpf). When exposed to ethanol from 10-18 hpf, *vangl2* and *gpc4* mutants had very similar variation in viscerocranial shape and size compared to mutant embryos treated from 10-24 hpf (Figure 3A-B). Interestingly, we did observe subtle differences in viscerocranial morphology between our *gpc4* mutant treatment groups. Unlike in Figure 2, where PC2 drove differences in viscerocranial shape between untreated and ethanol-treated *gpc4* mutants, exposure timing drove variation along PC1 (representing 87% of variation in data; Figure 3B). The mean shape shifted further left (negative for PC1) as the duration ethanol exposure increased. To test that ethanol interacts with *vangl2* and *gpc4* during C&E or in the transition between C&E and APE/CNC morphogenesis (prior to APE/CNC morphogenesis), we treated our fish with ethanol from either 6-10 hpf or 8-12 hpf and analyzed viscerocranial shape (Supplemental Figures 1 and 2, respectively). Neither mutant displayed any significant difference viscerocranial morphology between untreated and ethanol-treated mutants, when exposed to ethanol from 6-10 hpf or 8-12 hpf. To examine if ethanol interacts with *vangl2* and *gpc4* after APE/CNC morphogenesis, when the CNC have condensed into the pharyngeal arches, we treated mutant larvae with ethanol from 24-30 hpf. We again did not observe any significant ethanol-induced changes in viscerocranial shape (Supplemental Figure 3). Overall, this suggests that the ethanol sensitive time window for both *vangl2* and *gpc4* is between 10-18 hpf, after C&E and during APE/CNC morphogenesis.

**Figure 3.**
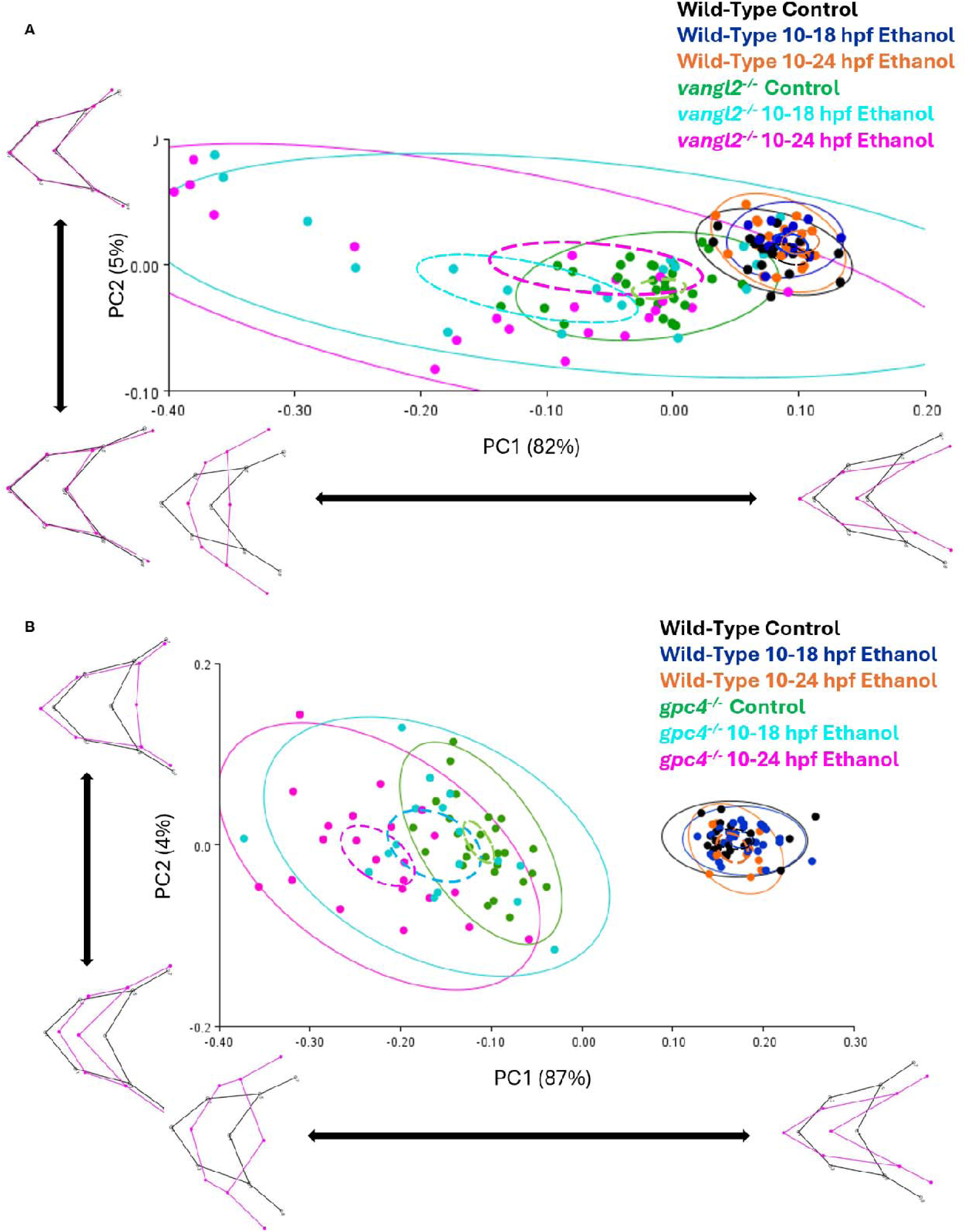
*vangl2* and *gpc4* mutant embryos are most sensitive to ethanol induced jaw defects between 10-18 hpf. **(A)** Principal component analysis plot and wireframes from morphometric analysis data. Landmarks were placed on the joints between the cartilage structures of the viscerocranium. Each genotype is color-coded: black = wild type control embryos (n=21), blue = wild type 10-18 hpf ethanol treated embryos (n=14), orange = wild type 10-24 hpf ethanol treated embryos (n=22), green = *vangl2^−/−^* control embryos (n=30), cyan = *vangl2^−/−^* 10-18 hpf ethanol treated embryos (n=22), magenta = *vangl2^−/−^* 10-24 hpf ethanol treated embryos (n=22). Procrustes ANOVA showed significant change in the viscerocranial shape (F = 16.75, DF = 60, p = <.0001). **(B)** principal component analysis plot and wireframes from morphometric analysis data. Landmarks were placed on the joints between the cartilage structures of the viscerocranium. Each genotype is color-coded: black = wild type control embryos (n= 20), blue = wild type 10-18 hpf ethanol treated embryos (n= 29), orange = wild type 10-24 hpf ethanol treated embryos (n=13), green = *gpc4^−/−^*control embryos (n= 28), cyan = *gpc4^−/−^* 10-18 hpf ethanol treated embryos (n= 17), magenta = *gpc4^−/−^* 10-24 hpf ethanol treated embryos (n= 22). solid circles represent confidence ellipses in which 95% of all individual data points for each group lie, dashed circles represent 95% confidence ellipses for means. Wireframe graphs represent variation termed by each axis with black representing no variation and magenta representing variation relative to the black wireframe. For example, PC1 captures a shortening and widening in viscerocranial shape, while PC2 represents variation in midfacial width. Procrustes ANOVA showed significant change in the viscerocranial shape (F =107.01, DF = 60, p =<0.0001).

### PCP mutants genetically interact during facial development

Previous work from Mile et al. (2017) showed that *vangl2* and *gpc4* genetically interact during C&E leading to exacerbated endodermal defects over the single mutants (Miles et al., 2017). We sought out to establish whether *vangl2* & *gpc4* also genetically interact and are further sensitized to ethanol-induced facial defects. To test this, we generated *vangl2^+/−^; gpc4^+/−^* double heterozygous carriers and crossed them to produce *vangl2*; *gpc4* compound heterozygous (heterozygous for one gene, homozygous mutant of the second gene) and double homozygous mutant larvae. Again, all larvae were treated with 1% Ethanol (v/v) from 10-24 hpf as described above. We observed severe facial and eye field defects in ethanol-treated compound heterozygous and homozygous double mutant larvae (Figure 4A-J, Table 1). However, due to the severe morphological changes in our mutant larvae, we were not able to perform our morphometric analyses of shape on compound heterozygotes and homozygous double mutants. To simplify our measures, we quantified the length of the Mc and Ch cartilages, as well as the internal angle of the Ch cartilages (Figure 4N-P, S-T). We also measure eye field separation from lens to lens (Figure 4M, R). To account for variation in size due to ethanol exposure, we used overall head length in our linear measures (Figure 4Q) and report our Mc and Ch length measurements as Meckel’s-to-Head length (Figure 4N) and Ceratohyal-to-Head length (Figure 4O) ratios. Looking at head length measures alone ethanol did not impact head length in wild-type and *gpc4* mutant larvae (Supplemental Figure 4). We observed ethanol-induced decreases in head length in *vangl2* mutants. Homozygous double mutant larvae were unaffected (Supplemental Figure 4).

**Figure 4.**
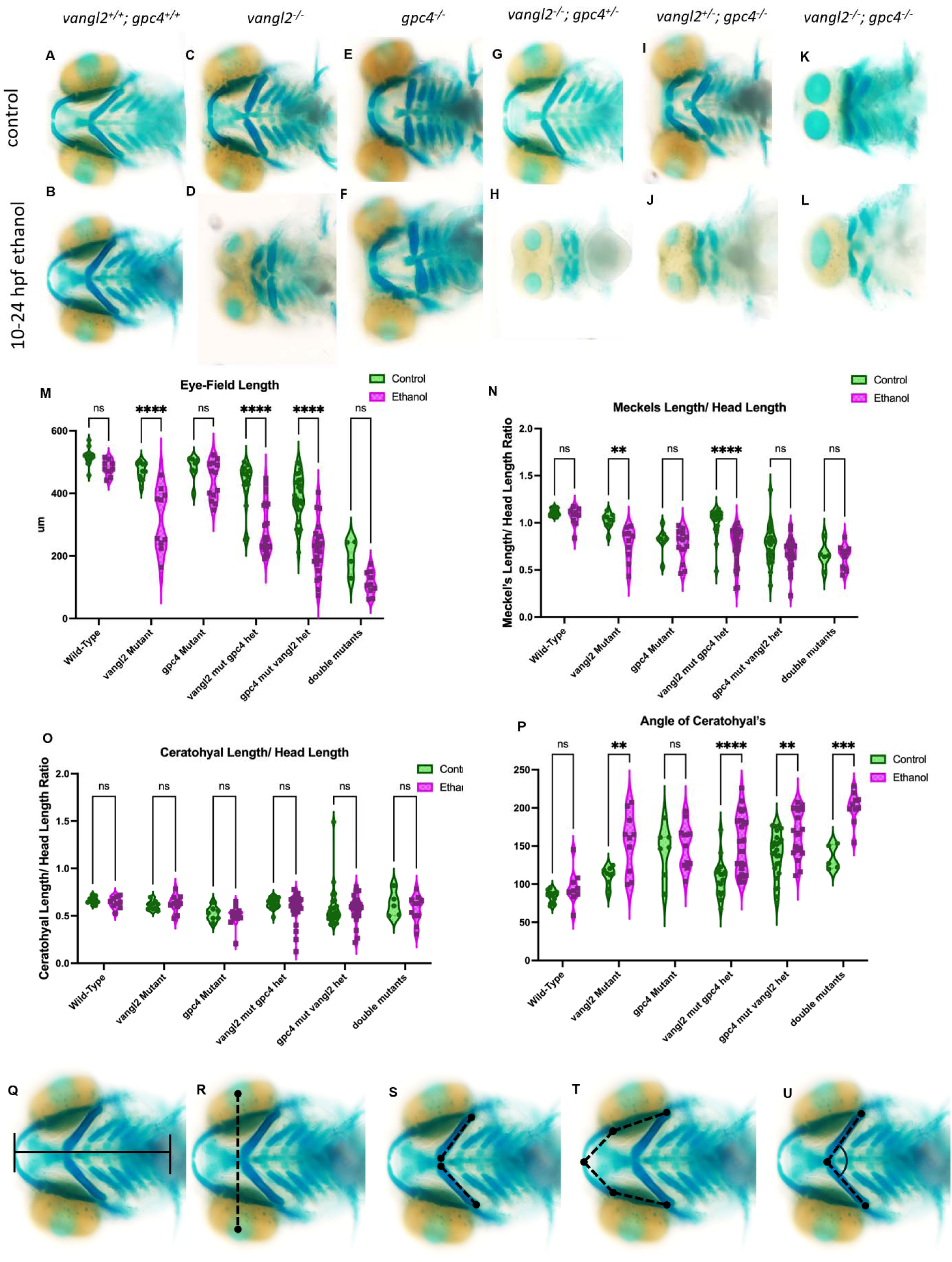
*vangl2* and *gpc4* genetically interact during jaw development. Whole-mount images of the viscerocranium of **(A)** wild type control embryos (n=11), **(B)** wild type ethanol treated embryos (n=9), **(C)** *vangl2^−/−^* control fish (n=9), **(D)** *vangl2^−/−^* ethanol treated fish (n=10), **(E)** *gpc4^−/−^* control fish (n=7), **(F)** *gpc4^−/−^*ethanol treated fish (n=13), **(G)** *vangl2^−/−^; gpc4^+/−^*control fish (n=16), **(H)** *vangl2^−/−^; gpc4^+/−^*ethanol treated fish (n=27), **(I)** *vangl2^+/−^; gpc4^−/−^* control fish (n=25), **(J)** *vangl2^+/−^; gpc4^−/−^* ethanol treated fish (n=23), **(K)** *vangl2^−/−^; gpc4^−/−^* control fish (n=5) and **(L)** *vangl2^−/−^; gpc4^−/−^* ethanol treated fish (n=11). **(M)** Eye-Field length. **(N)** Meckel length and **(O)** Ceratohyal length analyzed as ratios to overall head length. **(P)** Measures for the angle of the joints of the ceratohyals. Two-Way Anova was used to further analyze data. **(Q-U)** Whole-mount images of the viscerocranium in 4 dpf larvae showing linear measures and cartilage angles (cartilage is blue, ventral views, anterior to the left).

The ethanol-induced reduction in head length in larvae either heterozygous or homozygous mutant for *vangl2* was expected as the facial skeleton is significantly shorter and wider resulting in a shorter head length. Even after accounting for head length, we still observed significant ethanol-induced reductions in Mc length in *vangl2* single mutant larvae (Figure 4N). In our homozygous double mutants, we did not observe ethanol-induced significant changes in Mc cartilage lengths. This was not surprising given that untreated double mutants already displayed severe craniofacial defects (Figure 4K). Ethanol did not result in significant reductions in Mc length in *gpc4* mutants (Figure 4N). While, Ch length was unaffected in all groups (Figure 4N, O), ethanol did increase the size of the angle of Ch angle in our *vangl2* single mutants and homozygous double mutants (Figure 4P) compared to their untreated siblings.

Unlike our *vangl2* single mutants and homozygous double mutants, our compound heterozygous mutants mainly resemble their single mutant siblings when untreated (Figure 4G & I). However, when exposed to ethanol, many of our compound heterozygous larvae had exacerbated Mc and Ch cartilage defects and eye field defects, comparable to the double mutant larvae (Figure 4H, J compared to K, L). We observed that Mc length was significantly shorter in ethanol treated *vangl2* heterozygous compound mutant larvae (Figure 4N). There was also a significant ethanol-induced increase in Ch angle in *vangl2* heterozygous compound mutant larvae (Figure 4P). As with *gpc4* single mutants, there is no significant difference in Mc length *gpc4* compound mutant larvae although we do see a downward trend in that data (Figure 4N). We did observe a significant increase in Ch angle in ethanol-treated *gpc4* compound mutants (Figure 4P), demonstrating that a flattening, or even an inversion, of the Ch occurs. While we are not able to perform morphometric analysis on our compound heterozygous and homozygous double mutants, similar shape changes did occur, and they were reminiscent of our single mutants (Figures 1 & 2).

In addition to facial phenotypes, we observed increasing rates of ethanol-induced synophthalmia (with or without partial lens fusion) and cyclopia (full lens fusion) in our compound heterozygous and homozygous double mutants (Figure 4A-M, Table 1). As described above, we observed synophthalmia (without lens fusion) in ethanol-treated *vangl2* single mutants but not in *gpc4* single mutants (Figure 1, Table 1). Untreated homozygous double mutant larvae 80% showed synophthalmia (without lens fusion) and 20% cyclopia (Figure 4K, Table 1). Ethanol-treatment increased synophthalmia with partial lens fusions to 64% (from 0% untreated, Table 1), though we did observe a decrease in full cyclopia (20% to 9%, Table 1). This demonstrates that gpc4 loss potentiates facial shape defects in vangl2 mutants, independent of ethanol, but ethanol exacerbates these defects. In both untreated *gpc4* compound mutant larvae and *vangl2* compound mutant larvae, 88% had complete eye field separation while only 12% had synophthalmia without lens fusions (Table 1). However, ethanol treatment increased synophthalmia without lens fusions in *gpc4* compound mutant larvae and *vangl2* compound mutant larvae to 83% and 41%, respectively (Table 1). Strikingly, we observed 4% cyclopia in *gpc4* compound mutant larvae but 0% in *vangl2* compound mutant larvae. To quantify these eye defects, we measured the lens-to-lens width untreated and ethanol-treated single, compound heterozygous and homozygous double mutant larvae. Homozygous double mutant larvae did not display ethanol-induced reductions in lens-to-lens width, though we did see a downward trend (Figure 4M). This is largely due to severity already observed in untreated double mutants with downward trend representing increased lens fusions (Table 1). We observe significant ethanol-induced reductions in lens-to-lens width in *vangl2* single and compound heterozygous mutant larvae (Figure 4M). We also observed significant ethanol-induced reductions in lens-to-lens width in *gpc4* compound heterozygous mutants but not *gpc4* single mutants. This suggests that loss of one copy of *vangl2* sensitizes *gpc4* mutants to ethanol-induced synophthalmia. Taken together, our data shows that like our single mutants, our compound heterozygous and homozygous double mutants are ethanol sensitive after C&E and interact increasing the penetrance and expressivity of ethanol-induced facial defects.

### Loss of BMP signaling leads to exacerbated craniofacial phenotypes in PCP mutants

We have previously shown that multiple members of the BMP pathway, including the ligand *bmp4*, are ethanol sensitive from 10-18 hpf, disrupting endoderm morphogenesis and resulting in defects to the viscerocranium (Klem et al., 2025). Based on phenotypic and exposure paradigm similarities, we predicted that the BMP and PCP interact and sensitize embryos to ethanol-induced viscerocranial defects. To test this, we generated compound *bmp4; vangl2* and *bmp4; gpc4* double heterozygous crosses to analyze this potential interaction. Loss of one or both *bmp4* alleles further sensitize *vangl2* and *gpc4* mutants to ethanol, increasing both the penetrance and expressivity of jaw defects and synophthalmia (Figure 5A,B,D,G, Tables 2 & 3). We observed increases in severity of jaw hypoplasia, increases in synophthalmia in *vangl2* mutant / *bmp4* heterozygous larvae and homozygous double mutant larvae but not in *bmp4* mutant / *vangl2* heterozygous larvae (Table 2). We observed similar increases in *gpc4* mutant / *bmp4* heterozygous larvae and homozygous double mutant larvae only, including synophthalmia which has not been observed in *gpc4* single mutants (Table 3). Strikingly, we also observed several new craniofacial defects not originally seen in ethanol-treated BMP mutants or PCP mutants alone. We observed ectopic cartilages (Figure 5C), midline defects (which includes asymmetry of the midline joint between the ceratohyals; Figure 5E & F), basihyal defects (the basihyal is either split, crooked and/or rodded; Figure 5H, I & J) or palate defects (rodded ethmoid palate or jagged trabecula; Figure 5K & L). In our screen, these novel midline defects only marginally increased in ethanol suggesting they may be a novel ethanol-independent set of phenotypes in BMP/PCP interactions (Tables 2 & 3; Supplemental Figure 5). Taken together, our data show that disruptions to BMP signaling sensitizes PCP mutants and generates novel midline defects not observed in BMP or PCP mutants. This suggests that BMP/PCP interactions may lie at the heart of ethanol-sensitive endoderm morphogenesis driving facial defects.

**Figure 5.**
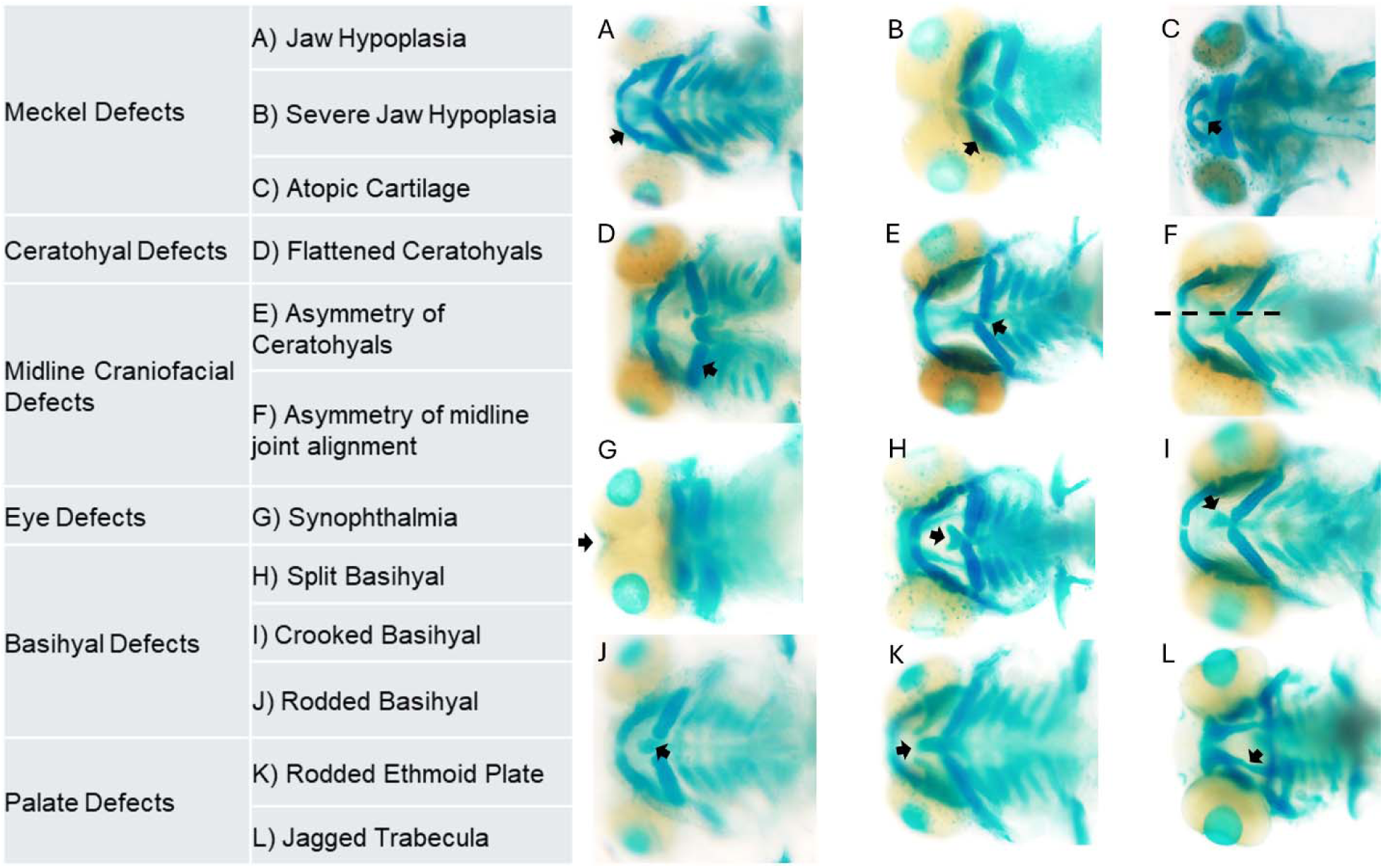
Representative images of phenotypes found in PCP/BMP crossed embryos. Whole-mount images of the malformations found in the viscerocranium of PCP/BMP embryos (ventral views, anterior to the left). the phenotypes displayed are **(A)** jaw hypoplasia, **(B)** severe jaw hypoplasia, **(C)** ectopic cartilage, **(D)** flattened ceratohyals, **(E)** asymmetry of ceratohyals, **(F)** asymmetry of midline joint alignment, **(G)** synophthalmia, **(H)** split basihyal, **(I)** crooked basihyal, **(J)** rodded basihyal, **(K)** rodded ethmoid plate and **(l)** jagged trabecula. arrows indicate the malformation or phenotype described.

**Table 2.**
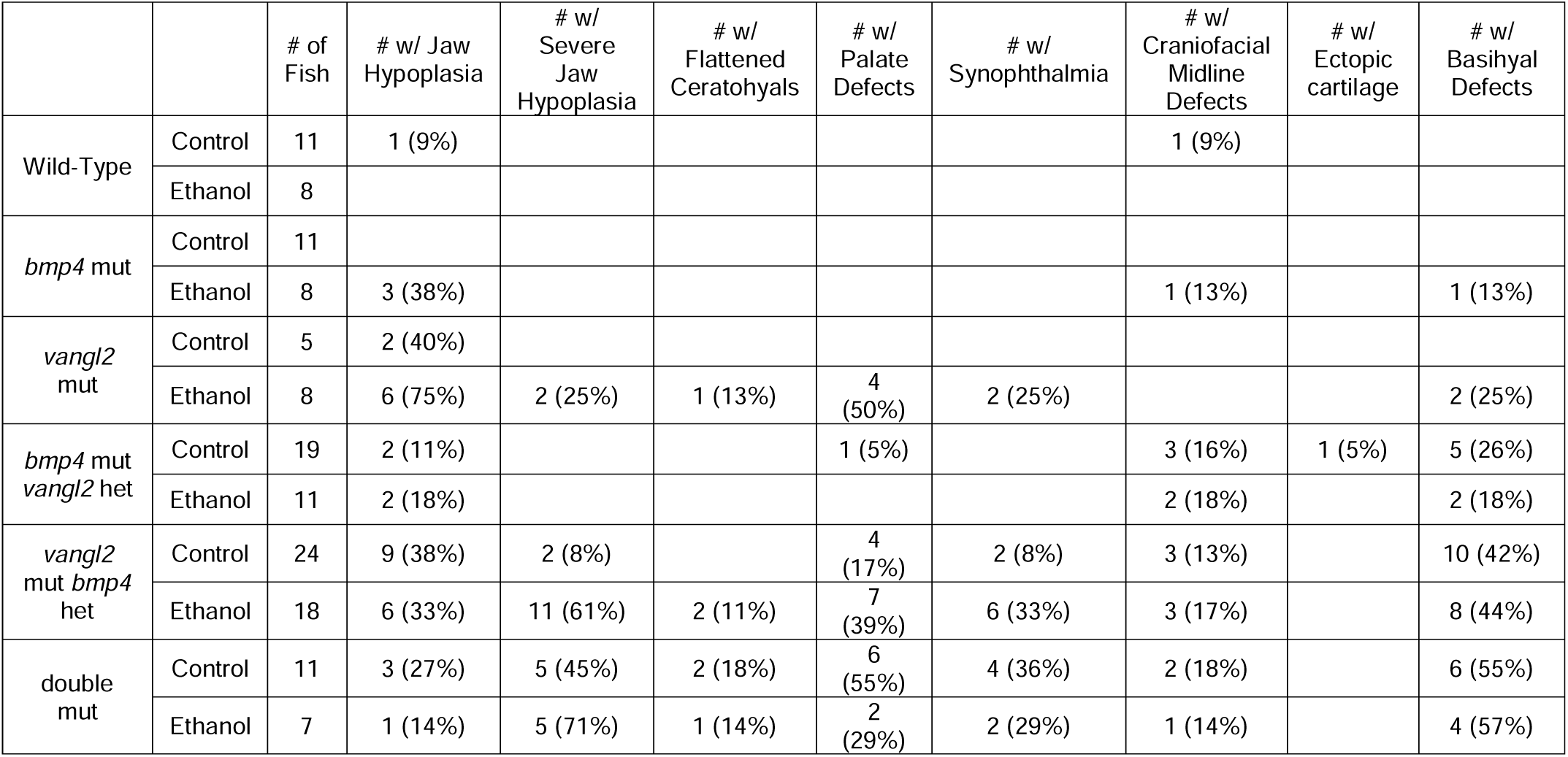
Summarized data on the occurrence of craniofacial phenotypes under different treatment conditions in *vangl2/bmp4* cross. Column A represents the genotype of the embryos (Wild-Type, *bmp4* mut, *vangl2* mut, *bmp4* mut *vangl2* het, *vangl2* mut *bmp4* het and double mut) while Column B defines the treatment condition (control or ethanol). Column C displays the total number of fish for each genotype under the specified conditions. Column D represents the number of fish in the group with jaw hypoplasia, and Column E represents the number of fish in the group with jaw loss. Jaw hypoplasia and jaw loss are mutually exclusive, meaning that if an embryo has one, it cannot have the other. Column F shows the number of fish in the group with ectopic cartilage, while Column G represents the number of fish with flattened ceratohyals. Column H provides the number of fish with craniofacial midline defects, Column I indicates the number with synophthalmia, and Column J details the number with basihyal defects. Column K represents the number of fish in the group with palate defects. The numerical values represent the number of fish that have the phenotype described within a given genotype and treatment group. Empty cells represent 0 phenotype observed in the group. The parentheses represent the percentage of embryos with the phenotype out of the total number of fish.

**Table 3.** Summarized data on the occurrence of craniofacial phenotypes under different treatment conditions in *gpc4/bmp4* cross. Column A represents the genotype of the embryos (Wild-Type, *bmp4* mut, *gpc4* mut, *bmp4* mut *gpc4* het, *gpc4* mut *bmp4* het and double mut) while Column B defines the treatment condition (control or ethanol). Column C displays the total number of fish for each genotype under the specified conditions. Column D represents the number of fish in the group with jaw hypoplasia, and Column E represents the number of fish in the group with jaw loss. Jaw hypoplasia and jaw loss are mutually exclusive, meaning that if an embryo has one, it cannot have the other. Column F shows the number of fish in the group with ectopic cartilage, while Column G represents the number of fish with flattened ceratohyals. Column H provides the number of fish with craniofacial midline defects, Column I indicates the number with synophthalmia, and Column J details the number with basihyal defects. Column K represents the number of fish in the group with palate defects. The numerical values represent the number of fish that have the phenotype described within a given genotype and treatment group. Empty cells represent 0 phenotype observed in the group. The parentheses represent the percentage of embryos with the phenotype out of the total number of fish.

|  |  | # of Fish | # w/ Jaw Hypoplasia | # w/ Severe Jaw Hypoplasia | # w/ Flattened Ceratohyals | # w/ Palate Defects | # w/ Synophthalmia | # w/ Craniofacial Midline Defects | # w/ Ectopic cartilage | # w/ Basihyal Defects |
| --- | --- | --- | --- | --- | --- | --- | --- | --- | --- | --- |
| Wild-Type | Control | 5 |  |  |  |  |  |  |  |  |
|  | Ethanol | 9 |  |  |  |  |  | 1 (11%) |  |  |
| *bmp4* mut | Control | 3 |  |  |  |  |  |  |  |  |
|  | Ethanol | 6 | 1 (17%) |  |  |  |  |  |  | 2 (33%) |
| *gpc4* mut | Control | 9 | 7 (78%) | 2 (22%) |  | 4 (44%) |  | 1 (11%) |  | 2 (22%) |
|  | Ethanol | 6 | 2 (33%) | 4 (67%) | 1 (17%) | 3 (50%) |  | 1 (17%) | 3 (50%) | 2 (33%) |
| *bmp4* mut *gpc4* het | Control | 21 | 3 (14%) |  |  |  |  | 3 (14%) |  | 1 (5%) |
|  | Ethanol | 10 |  | 1 (10%) |  |  |  | 4 (40%) |  |  |
| *gpc4* mut *bmp4* het | Control | 12 | 11 (92%) | 1 (8%) | 1 (8%) | 5 (42%) |  | 1 (8%) |  | 5 (42%) |
|  | Ethanol | 17 | 5 (29%) | 12 (71%) | 7 (41%) | 12 (71%) | 1 (6%) | 3 (18%) | 3 (18%) | 8 (47%) |
| double mut | Control | 4 | 3 (75%) | 1 (25%) | 2 (50%) | 3 (75%) |  | 1 (25%) | 1 (25%) | 3 (75%) |
|  | Ethanol | 16 | 3 (19%) | 13 (81%) | 5 (31%) | 5 (31%) | 2 (13%) | 3 (19%) | 3 (19%) | 6 (38%) |

## Discussion

Prenatal alcohol exposure (PAE) results in a wide range of structural malformations, including those to the facial skeleton (Sokol et al. 2003; Hoyme et al. 2016). Described as Fetal Alcohol Spectrum Disorders (FASD), the malformations include, thin upper lip, microcephaly and, for purposes of this work, jaw hypoplasia (Lovely, 2020). Formation of the facial skeleton is a highly complex process requiring reciprocal signaling events between multiple cell types, including the pharyngeal endoderm and the cranial neural crest (CNC). These signaling interactions rely greatly on separate morphogenesis events in each tissue (Swartz et al., 2012; Lovely et al., 2016). We and others have shown that many of the genetic loci that regulate these events can be sensitive to ethanol, disrupting facial formation (Swartz et al., 2014; Klem et al., 2025; Vo and Lovely, 2025). One such set of ethanol-sensitive genetic pathways is the highly conserved Planar Cell Polarity (PCP) pathway. PCP plays a major role in a host of developmental processes, including convergence and extension (C&E), endoderm morphogenesis and facial development (Miles et al., 2017; Scar et al., 2021; Ulrich et al., 2003; Ulrich et al., 2005, Ling et al., 2017). Loss of either *vangl2* or *gpc4* results in C&E defects in the endoderm leading to a shorter trunk and wider body axis, which is exacerbated in *vangl2; gpc4* double mutants (Miles et al., 2017). Moreover, *gpc4* has been shown to play a role in cartilage polarity and cell stacking during neural crest development (Sisson et al., 2015; Ling et al., 2017).

### PCP-ethanol interactions lead to facial defects

Previous work showed that *vangl2* and *gpc4* mutants are sensitive to ethanol-induced facial defects and synophthalmia when treated from 6-24 hpf (Sidik et al., 2021). However, this time window covers both C&E (6-10 hpf) and endoderm/CNC morphogenesis (10-24 hpf). This work examining ethanol-induced phenotypes in *vangl2* and *gpc4* mutants never separated these two developmental events through varying the time windows of ethanol exposure. To focus on the ethanol-sensitive role that *vangl2* and *gpc4* play in endoderm/CNC morphogenesis, we examined multiple ethanol-exposure time windows spanning C&E (6-10 hpf), endoderm/CNC morphogenesis (10 hpf-24 hpf) and after the CNC condense into the pharyngeal arches (24-30 hpf). We observed that both *vangl2* and *gpc4* are ethanol sensitive during endoderm/CNC morphogenesis (10-24 hpf), but not C&E (6-10 hpf). In addition, both mutants were insensitive to ethanol after 24 hpf demonstrating that *vangl2* and *gpc4* were ethanol sensitive during endoderm/CNC morphogenesis (10-24 hpf) leading to increased synophthalmia and facial defects. However, given that our focus was on facial development, we did not examine expression changes in C&E markers that Sidik et al., (2021) observed in their work. It is possible that when isolated, the significant reductions in expression domain size of these C&E markers may not manifest in the subsequent facial phenotypes. This suggests that the facial phenotypes we observed in our 10-24 hpf exposure window result from additional developmental events occurring after C&E, in particular endoderm/CNC morphogenesis. Future work varying exposure paradigms and broader mechanistic analyses will be needed to fully address this discrepancy.

Further analysis with our morphometric approach showed subtle variation in facial shape that was not previously observed. Ethanol-treated wild-type larvae displayed altered facial morphology compared to untreated wild-type siblings, though the shape change was not as pronounced as observed in untreated *vangl2* mutants. This suggests ethanol increases facial variation and loss of *vangl2* potentiates these defects. While we did not observe similar differences between untreated and ethanol-treated *gpc4* mutants when exposed from 10-24 hpf, we did see that when exposed from 10-18 hpf the facial shape changes were not as expressive as those exposed 10-24 hpf. Given the *gpc4* plays roles in both endoderm morphogenesis and CNC development (Sisson et al., 2015; Miles et al., 2017; Ling et al., 2017), the subtle differences in facial shape in the different exposure windows are not surprising. It is possible that *gpc4* may be playing roles in two different yet mechanistically linked developmental processes driving facial development. While our exposure time windows suggest these distinct temporal roles of *gpc4*, future will be needed to separate these dual roles for *gpc4* in facial development. Overall, this suggests that while exposure and genotype drive facial shape changes in *vangl2* mutants, timing of exposure drives these changes in *gpc4* mutants and *vangl2* mutation fully sensitizes larvae to ethanol-induced facial defects but *gpc4* only mildly sensitizes larvae depending on the exposure window.

While *vangl2*- or *gpc4*-ethanol interactions result in unique facial malformations, they both impact facial development suggesting they interact hyper-sensitizing larvae to ethanol. Here, we show that heterozygous loss of *gpc4* sensitized *vangl2* larvae to ethanol-induced facial malformations and increased synophthalmia/cyclopia, consistent with the work from Sidik et al., (2021). However, our work narrows the *vangl2*- and *gpc4*-ethanol interactions to endoderm/CNC morphogenesis after C&E (10-24 hpf), suggesting that either CNC development or endoderm morphogenesis (or both) may be impacted by these interactions. Previous work has shown that *gpc4* is necessary for CNC development in part by regulating CNC polarity (Ling et al., 2017). However, loss of *gpc4* does not appear to disrupt pharyngeal arch morphology at 36 hpf though it does act cell autonomously in the CNC suggesting it plays a role during CNC morphogenesis out of the pharyngeal arches to form the facial skeleton (Sisson, et al., 2015). While it’s possible that ethanol may be interacting with *gpc4* mutants to exacerbate polarity defects earlier during endoderm/CNC morphogenesis that then result in later CNC developmental defects, we can’t rule out that ethanol may be synergizing with *gpc4* mutants through additional targets. It is possible that *vangl2* is the target of interaction with ethanol as we see *vangl2* mutation sensitizing larvae to ethanol-induced facial malformations. This suggests that CNC develop could be disrupted in ethanol-treated *vangl2* mutants.

While *vangl2* mutants do impact CNC migration, this impact is mild and may not fully explain the impact of ethanol on facial development in *vangl2* mutants (Matthews et al., 2008). However, ethanol does exacerbate facial phenotypes in mutants of CNC migration (McCarthy et al., 2013). This suggests that *vangl2* mutation may sensitize CNC to ethanol-induced defects not observed in ethanol or *vangl2* loss of function alone. For *gpc4* single mutants, ethanol may be disrupting *vangl2* function, impacting CNC development. It is also possible that *vangl2* and *gpc4* may be disrupting endoderm morphogenesis. Both play roles in endoderm C&E (Miles et al., 2017) and cannot be ruled out of impacting endoderm morphogenesis after C&E and disrupting endoderm signaling to the CNC resulting in facial malformations. This would be consistent with our previous results showing that BMP-ethanol interactions disrupt endoderm morphogenesis driving CNC defects (Klem et al., 2025; Vo and Lovely, 2025). While we have previously shown that BMP signaling is required in the endoderm for its morphogenesis (Lovely et al., 2016), we would need to test cell autonomy of *vangl2* and *gpc4* to determine whether morphogenesis of the endoderm, CNC, both or neither are impacted in our ethanol-treated mutants.

### BMP-PCP interactions yield novel phenotypes

We have previously shown that multiple BMP mutants are ethanol-sensitive from 10-18 hpf leading to disrupted endoderm morphology (Klem et al., 2025). However, ethanol does not attenuate BMP signaling nor its downstream target *nkx2.3* but does increase CNC apoptosis in BMP mutants through a yet unknown signaling mechanism (Klem et al., 2025; Vo and Lovely, 2025). Throughout our study, we see overlapping similarities between the BMP and PCP pathways. BMP and PCP mutant larvae share ethanol sensitive time window of 10-18 hpf during endoderm/CNC morphogenesis (Klem et al., 2025). Ethanol-treated PCP mutants phenocopy ethanol-induced shorter and wider facial shape observed in BMP mutants (Klem et al., 2025). Both BMP and PCP regulate endoderm development (Lovely et al., 2016; Miles et al., 2017). All this strongly supports our analysis of BMP-PCP interactions in ethanol-sensitive facial development where we observed that loss one or both *bmp4* alleles further sensitizes PCP mutants to ethanol-induced facial defects and synophthalmia, phenocopying *vangl2; gpc4* double mutants and showing increased penetrance of facial defects and synophthalmia. Surprisingly, we observed additional midline craniofacial defects in untreated larvae not seen in our PCP double mutants. Here, ethanol mildly increases the penetrance of these midline defects. This suggests that the PCP and BMP pathways interact in two ways: 1. An ethanol-dependent exacerbation facial defects and synophthalmia; 2. An ethanol-independent generation of novel midline defects. Given the complexity of facial development, several mechanisms may be driving these two distinct interactions.

BMP may be upstream, regulating PCP component gene expression in the endoderm. This would lead to defects in endoderm morphogenesis resulting in facial defects. However, this does not explain how we only see synophthalmia in PCP and BMP-PCP mutants but not BMP mutants alone. It also leaves the open question of how defects in endoderm morphogenesis give rise to CNC apoptosis. It is possible that BMP drives ethanol-induced defects in endoderm morphogenesis while PCP drives ethanol-induced defects in the CNC. Here we would predict a synergy between BMP-endoderm and PCP-CNC defects driving the overall phenotypes. If this were true, we would predict that increased CNC apoptosis in ethanol-treated PCP mutants would result in similar facial malformations as seen in ethanol-treated BMP mutants (Klem et al., 2025). However, our morphometric and genetic interaction analyses argue against this as PCP mutants, both single and double, do not fully recapitulate the ethanol-induced facial phenotypes of BMP mutants. While ethanol-treated PCP mutants do not fully phenocopy ethanol-treated BMP mutants, it is possible that ethanol-induced CNC apoptosis in PCP mutants leads to unique outcomes in CNC development compared to ethanol-treated BMP mutants. Building on this, the novel midline phenotypes in our PCP/BMP double mutants suggest a more complex relationship between BMP and PCP signaling. We previously showed that BMP signaling regulates formation of lateral endoderm (Lovely et al., 2016) and PCP plays a major role in cell endoderm cell polarity (Miles et al., 2017). The presence of midline defects in our PCP/BMP double mutants suggests that medial-lateral patterning may also be affected leading to exacerbation of existing phenotypes and generation of the novel midline defects. Beyond studies of cell autonomy described above, detail analyses of BMP and PCP signaling dynamics and cell behavior will be needed to dissect the complex ethanol-sensitive relationship between BMP and PCP signaling.

Combined our results show that work in animal models like zebrafish are key to dissecting the complex relationship between genetics and ethanol exposure paradigms driving ethanol-induced development defects. Given the genetic and developmental homology between zebrafish and humans, our work provides a powerful model to examine complex genetic signaling pathways and generate a deeper mechanistic understanding of ethanol interactions with these pathways on craniofacial development. Here, we show that PCP mutants are ethanol sensitive after C&E, during endoderm/CNC morphogenesis, and interact with BMP signaling resulting in unique facial defects and synophthalmia. The expressivity and penetrance of these phenotypes suggest a novel and complex ethanol-sensitive signaling pathway driving endoderm/CNC morphogenesis during facial development. Future studies to identify the mechanism on a cellular level will be needed to dissect the role PCP and BMP during endoderm/CNC morphogenesis, and how ethanol fits into the story. Ultimately, this will continue to build on a conceptual framework and a mechanistic paradigm of ethanol-induced birth defects we have been developing, connecting ethanol exposure with concrete cellular events that could be sensitive beyond facial development.

---

**Supplemental Figure 1.**
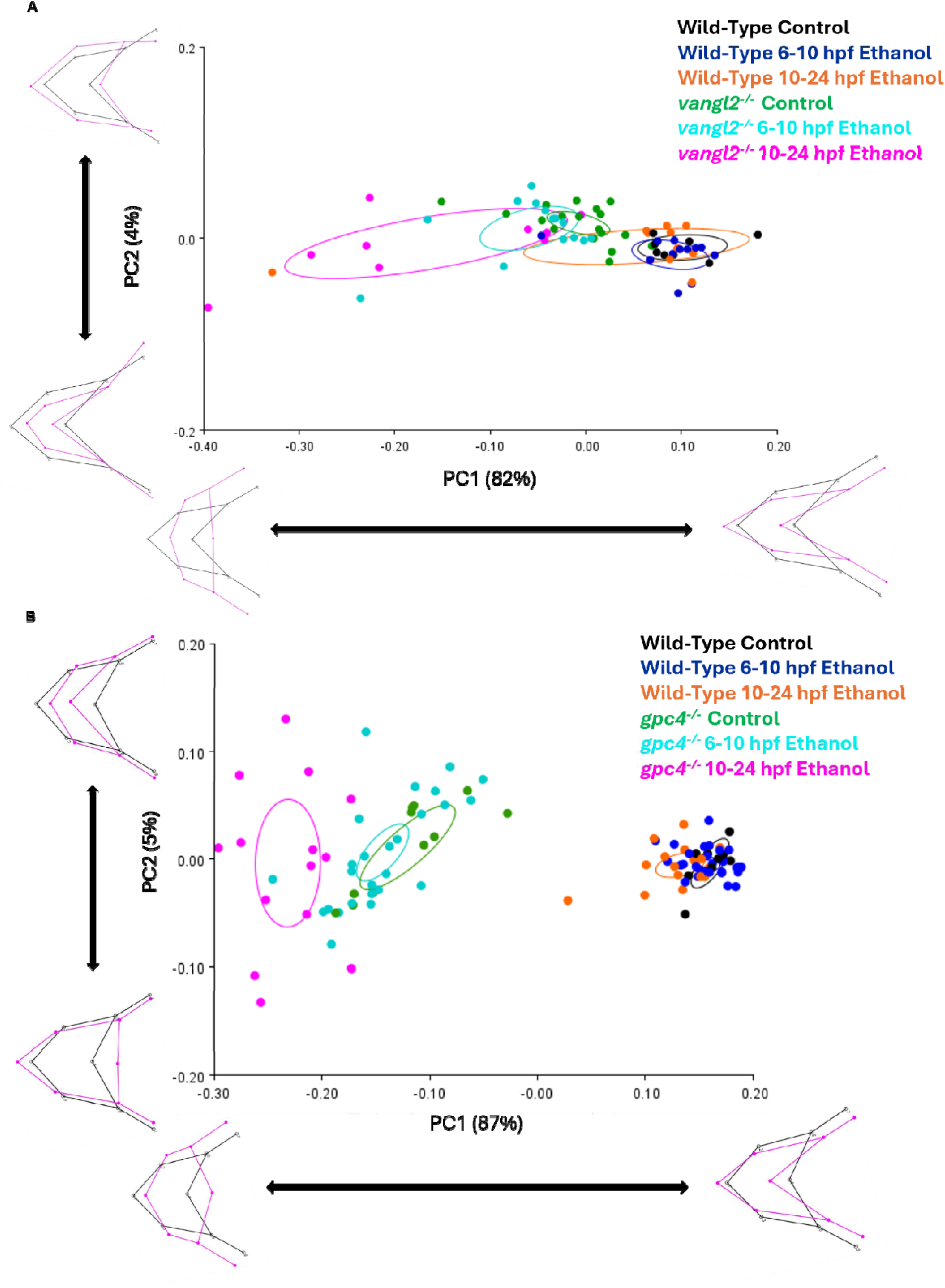
*vangl2* and *gpc4* mutant embryos are not sensitive to ethanol induced jaw defects when treated between 6-10 hpf. **(A)** principal component analysis plot and wireframes from morphometric analysis data. Landmarks were placed on the joints between the cartilage structures of the viscerocranium. Each genotype is color-coded: black = wild type control embryos (n=8), blue = wild type 6-10 hpf ethanol treated embryos (n=12), orange = wild type 10-24 hpf ethanol treated embryos (n=11), green = *vangl2^−/−^* control embryos (n=18), cyan = *vangl2^−/−^* 6-10 hpf ethanol treated embryos (n=14), magenta = *vangl2^−/−^* 10-24 hpf ethanol treated embryos (n=13). Solid circles represent 95% confidence ellipses for means. Procrustes ANOVA showed significant change in the viscerocranial shape (F = 11.34, DF = 60, p = <.0001). **(B)** principal component analysis plot and wireframes from morphometric analysis data. Landmarks were placed on the joints between the cartilage structures of the viscerocranium. Each genotype is color-coded: black = wild type control embryos (n= 9), blue = wild type 6-10 hpf ethanol treated embryos (n= 28), orange = wild type 10-24 hpf ethanol treated embryos (n=18), green = *gpc4^−/−^*control embryos (n= 11), cyan = *gpc4^−/−^* 6-10 hpf ethanol treated embryos (n= 26), magenta = *gpc4^−/−^* 10-24 hpf ethanol treated embryos (n= 14). Solid circles represent 95% confidence ellipses for means. Procrustes ANOVA showed significant change in the viscerocranial shape (F =102.67, DF = 60, p =<0.0001).

**Supplemental Figure 2.**
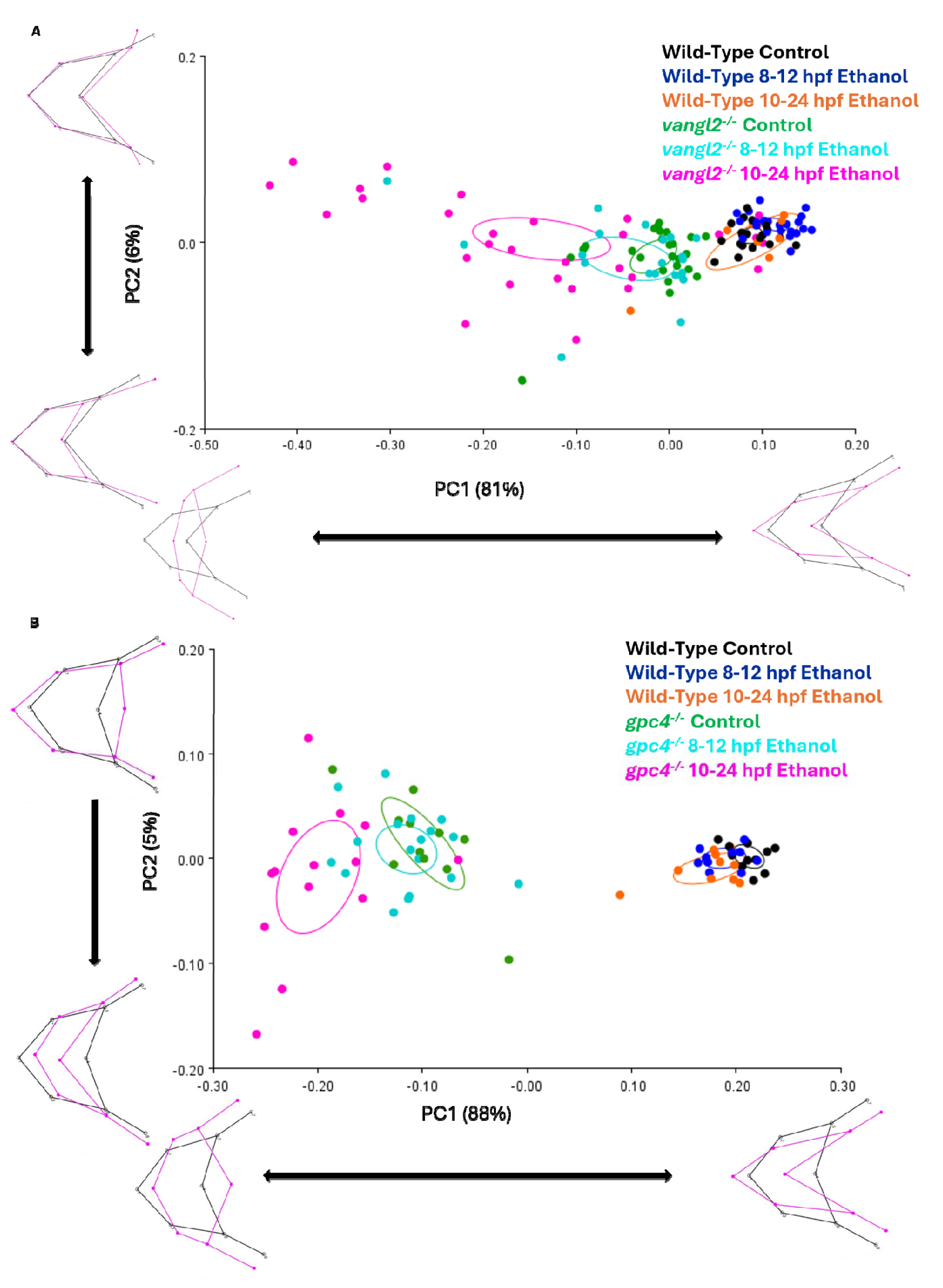
*vangl2* and *gpc4* mutant embryos are not sensitive to ethanol induced jaw defects when treated between 8-12 hpf. **(A)** principal component analysis plot and wireframes from morphometric analysis data. Landmarks were placed on the joints between the cartilage structures of the viscerocranium. Each genotype is color-coded: black = wild type control embryos (n=21), blue = wild type 8-12 hpf ethanol treated embryos (n=23), orange = wild type 10-24 hpf ethanol treated embryos (n=10), green = *vangl2^−/−^* control embryos (n=25), cyan = *vangl2^−/−^* 8-12 hpf ethanol treated embryos (n=21), magenta = *vangl2^−/−^* 10-24 hpf ethanol treated embryos (n=30). Solid circles represent 95% confidence ellipses for means. Procrustes ANOVA showed significant change in the viscerocranial shape (F = 22.28, DF = 60, p = <.0001). **(B)** principal component analysis plot and wireframes from morphometric analysis data. Landmarks were placed on the joints between the cartilage structures of the viscerocranium. Each genotype is color-coded: black = wild type control embryos (n= 11), blue = wild type 8-12 hpf ethanol treated embryos (n= 10), orange = wild type 10-24 hpf ethanol treated embryos (n=10), green = *gpc4^−/−^*control embryos (n= 11), cyan = *gpc4^−/−^* 8-12 hpf ethanol treated embryos (n= 18), magenta = *gpc4^−/−^* 10-24 hpf ethanol treated embryos (n= 14). Solid circles represent 95% confidence ellipses for means. Procrustes ANOVA showed significant change in the viscerocranial shape (F =73.99, DF = 60, p =<0.0001).

**Supplemental Figure 3.**
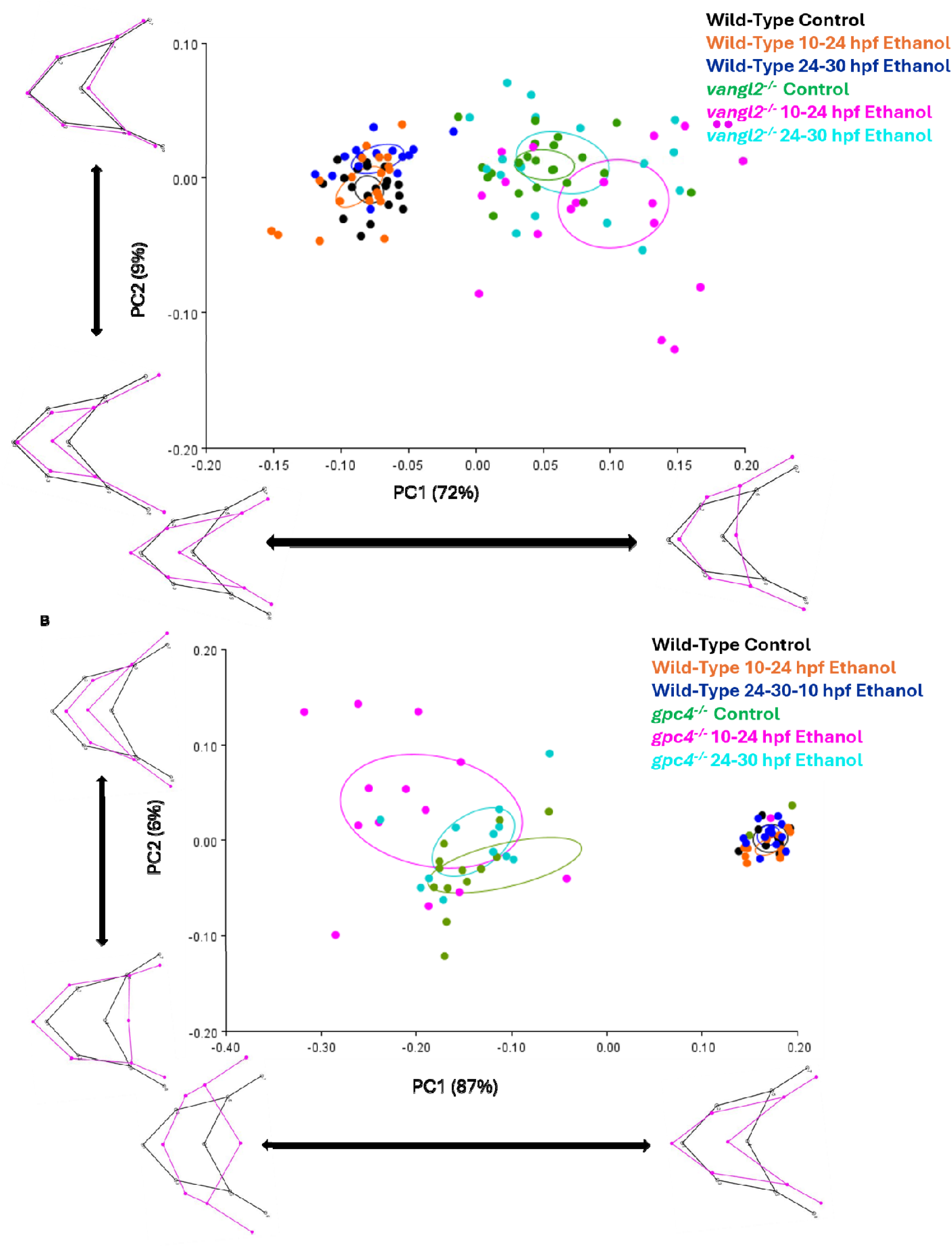
*vangl2* and *gpc4* mutant embryos are not sensitive to ethanol induced jaw defects when treated between 24-30 hpf. **(A)** principal component analysis plot and wireframes from morphometric analysis data. Landmarks were placed on the joints between the cartilage structures of the viscerocranium. Each genotype is color-coded: black = wild type control embryos (n=19), blue = wild type 24-30 hpf ethanol treated embryos (n=15), orange = wild type 10-24 hpf ethanol treated embryos (n=19), green = *vangl2^−/−^*control embryos (n=22), cyan = *vangl2^−/−^*24-30 hpf ethanol treated embryos (n=18), magenta = *vangl2^−/−^*10-24 hpf ethanol treated embryos (n=19). Solid circles represent 95% confidence ellipses for means. Procrustes ANOVA showed significant change in the viscerocranial shape (F = 30.51, DF = 60, p = <.0001). **(B)** principal component analysis plot and wireframes from morphometric analysis data. Landmarks were placed on the joints between the cartilage structures of the viscerocranium. Each genotype is color-coded: black = wild type control embryos (n= 9), blue = wild type 24-30 hpf ethanol treated embryos (n= 13), orange = wild type 10-24 hpf ethanol treated embryos (n=13), green = *gpc4^−/−^* control embryos (n= 16), cyan = *gpc4^−/−^*24-30 hpf ethanol treated embryos (n= 12), magenta = *gpc4^−/−^* 10-24 hpf ethanol treated embryos (n= 14). Solid circles represent 95% confidence ellipses for means. Procrustes ANOVA showed significant change in the viscerocranial shape (F =37.92, DF = 60, p =<0.0001).

**Supplemental Figure 4.**
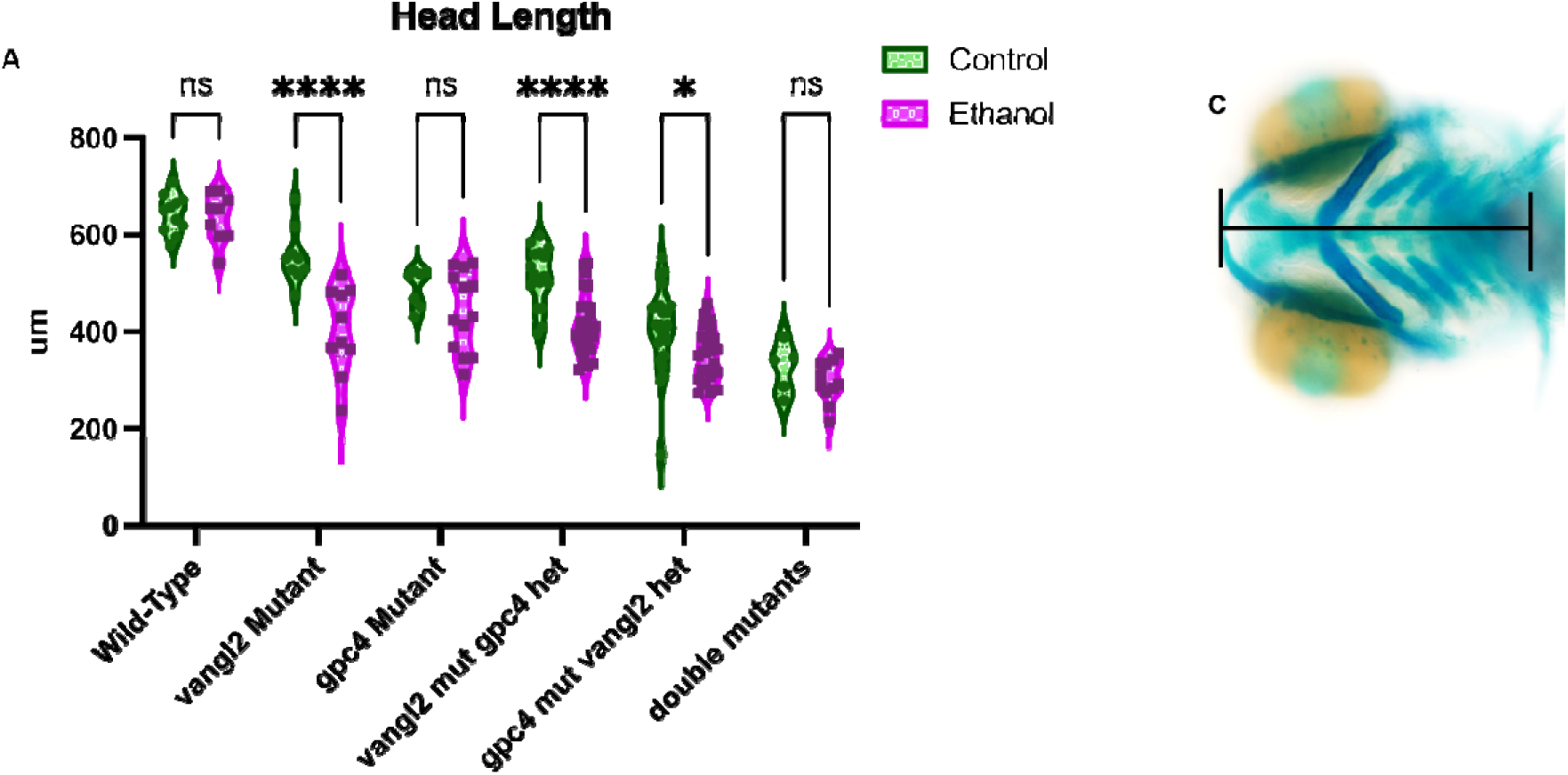
Comparison of Head length across all genotypes. Wild-type control embryos (n=11), Wild-Type ethanol treated embryos (n=9), *vangl2^−/−^* control fish (n=9), *vangl2^−/−^*ethanol treated fish (n=10), *gpc4^−/−^* control fish (n=7), *gpc4^−/−^*ethanol treated fish (n=13), *vangl2^−/−^; gpc4^+/−^* control fish (n=16), *vangl2^−/−^; gpc4^+/−^* ethanol treated fish (n=27), *vangl2^+/−^; gpc4^−/−^* control fish (n=25), *vangl2^+/−^; gpc4^−/−^* ethanol treated fish (n=23), *vangl2^−/−^; gpc4^−/−^* control fish (n=5) and *vangl2^−/−^; gpc4^−/−^* ethanol treated fish (n=11) were the genotypes used. **(A)** Comparison of head lengths of ethanol treated and untreated embryos within genotypes groups. Two-Way Anova was used to further analyze data.

**Supplemental Figure 5.**
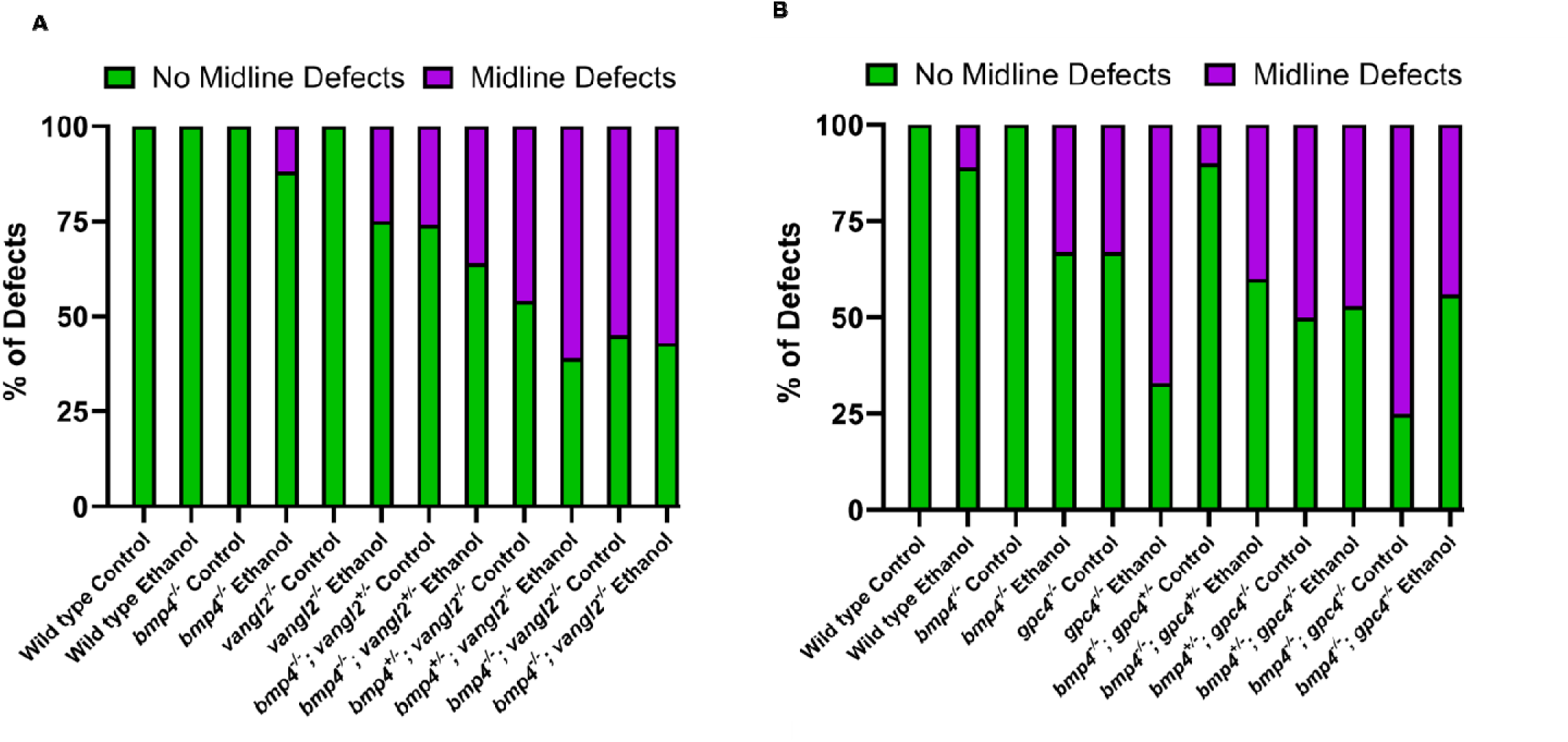
Bar graphs demonstrate the occurrence of midline defects in PCP/BMP mutant crosses. A) The occurrence of midline defects in *bmp4^+/−^; vangl2^+/−^*cross. B) The occurrence of midline defects in *bmp4^+/−^; gpc4^+/−^* cross. The percentages for the number of embryos that have no midline defects (Green) and those that have Midline defects (Magenta) are displayed for all genotype and treatment groups.

